# Role of the NHE1 exchanger in the antitumor effects of copper(II) complexes and phenanthroline derivatives

**DOI:** 10.1101/2024.12.21.629901

**Authors:** K.S. Muñoz Garzón, V. Martínez, V. de Giusti, D. Pérez Visñuk, M. Villaverde, N. Alvarez, G. Facchin, A.L. Di Virgilio

## Abstract

Three copper(II) complexes containing 1,10-phenanthroline ([CuCl_2_(phen)]·4H_2_O,**1**), neocuproine ([CuCl_2_(neo)]·4H_2_O, **2**) and tetramethyl-phenanthroline ([CuCl_2_(tmp)]·4H_2_O, **3**) as the primary ligand and another three copper(II) complexes with the L-Ala-Phe dipeptide as an auxiliary ligand: [Cu(L-Ala-Phe)(phen)]·4H_2_O (**4**), [Cu(L-Ala-Phe)(neo)]·4H_2_O (**5**) and [Cu(L-Ala-Phe)(tmp)]·4H_2_O (**6**), inhibited cell viability in breast cancer MCF-7 cell line, both in the monolayer and spheroid cell culture models. The pair with tetramethyl-phenanthroline displayed a better selectivity index than cisPt and non-cytotoxicity-related ROS induction and apoptosis in the monolayer breast cancer model. Cell proliferation was affected by all compounds in a concentration-dependent manner, with a more substantial effect on the tetramethyl-phenanthroline complexes. Cell viability on multicellular spheroids showed a concentration-dependent reduction from 1 μM, with IC_50_ that were half the one for cisplatin. All copper complexes, except for **1** showed DNA damage, demonstrated by the comet assay at a concentration below the IC_50_. The role of NHE1 has been linked to many types of cancers. Our study revealed that all compounds inhibited NHE1 activity in MCF-7 cells. However, only complexes containing the dipeptide auxiliary ligand could extend their effect on cell migration (Wound Healing Assay) and MMP-9 activity studied by zimography. Wester Blot analysis showed that expressions of MMP-2, MMP-9, and NHE1 were affected when MCF7 cells were treated with the six compounds as well. Overall, our results reveal an antitumor effect of all copper(II) complexes studied in breast cancer cells and a fundamental role of NHE1 in cell migration.

## 1. Introduction

Breast cancer is the second most diagnosed cancer in the world, with an incidence of 71.3% for both sexes in 2022 (1). Most breast cancers are of the luminal A subtype, described by estrogen receptor and progesterone receptor expression, but lacking the human epidermal growth factor receptor. Since they are estrogen-dependent cancers, they can be effectively treated by inhibiting estrogen receptor function. However, as the tumor progresses, it may develop resistance through various mechanisms, which is the cause of most breast cancer therapy failures, demanding novel chemotherapies for the evolving disease (2).

During these past decades, researchers have conducted multiple investigations to treat different types of cancer using metal complexes, including copper, demonstrating antitumor activities *in vitro* and *in vivo* (3–6) . Copper is a relevant trace element for normal physiological functions. Nevertheless, it can induce cell death at both low or high concentrations; this type of cell death is known as cuproptosis, occurs when copper accumulates within the mitochondria(7). Various copper complexes containing phenanthroline (phe) have been investigated. Back in 1979, [Cu(Phen)_2_]^2+^ was described as the first nuclease chemical agent (8). The nuclease activity induces DNA double-strand breaks through reactive oxygen species (ROS) formation, damaging and degrading DNA (9,10). The interaction with DNA makes copper(II) complexes good candidates for developing efficient antitumor treatment (11).

Neocuproine (neo=2,9-dimethyl-1,10-phenanthroline) is an ortho-substituted ligand first reported by Mohindru et al (12) and demonstrated to be cytotoxic in tumoral cells in a copper-dependent manner. Its ionophoric ability improves the mobility and interaction with cellular membranes (13) and increases cellular uptake and cytotoxicity by enhancing the lipophilicity of the complexes (14). On the other hand, studies have shown that 3,4,7,8-tetramethyl-1,10-phenanthroline (tmp) is highly cytotoxic to cancer cells effectively killing osteosarcoma cells at nanomolar concentrations with potency surprisingly better than cisplatin and carboplatin (15,16).

Amino acids are vital for building proteins and chemical species that perform several biological functions (17). L-phenylalanine is an essential amino acid to make melanin, dopamine, noradrenaline, adrenaline, and thyroxine. Its hydrophobicity makes it found buried within a protein (18). L-alanine is a nonessential amino acid uptaken by neutral amino acid transporters like Alanine, Serine, and Cysteine Transporter 2 (ASCT2) that transports glutamine for the biosynthesis of proteins (19). New studies have shown the critical role of amino acids in cancer metabolism (20). Cancer cells have been observed to use amino acids to increase cell proliferation through the induction of a signaling pathway that leads to glutamine transport inside cancer cells, biosynthesis of macromolecules, and generation of energy (19,21).

Moreover, some studies have investigated the therapeutic potential of dipeptides for breast cancer treatment. An endogenous dipeptide composed of β-alanine and L-histidine possesses anti-inflammatory and antioxidant activities. It modulates cell proliferation, cell cycle arrest, apoptosis, and glycolytic energy metabolism, decreasing cancer cells’ size and viability (22). Moreover, complexes containing phenanthroline and tyrosine inhibited the growth of breast cancer cells with less toxicity, inducing apoptosis and autophagy (23).

The sodium-hydrogen exchanger (NHE) is a transmembrane protein that facilitates the exchange of one proton (H^+^) for one sodium ion (Na^+^) across lipid bilayers, thereby regulating the intracellular pH and Na^+^ concentration. Among its isoforms, isoform 1 (NHE1) is the most important and, in addition to its fundamental role in pH and Na^+^ homeostasis, NHE1 has been shown to participate in cell cycle regulation, proliferation, migration and adhesion, and the promotion of apoptosis resistance(24). Tumor cells are characterized by extracellular acidosis due to the high proliferation rate and the expulsion of protons from the cytosol to the surrounding environment, causing acidification in the confined and poorly vascularized extracellular space. The resulting alkaline intracellular pH favors the proliferation (25) enhances glycolytic metabolism (26) and counteracts apoptosis (27), in contrast, an acidic extracellular pH drives tumor cell migration and invasion (28). Therefore, a malignant, highly invasive cancer cell phenotype is acquired in an acidic environment (29–31) and, in many cases, is induced by NHE1 (32). The role of NHE1 has been linked to many types of cancers such as breast tumors (33,34) among others. Efficient cell migration depends partly on the fact that NHE1 activity generates acidic intracellular pH that controls cell-matrix adhesion and the cytoskeletal migration machinery (35).

Remodeling the extracellular matrix, a crucial event in the migration process, is orchestrated by metalloproteases (MMPs). High levels of MMPs are distinctive in many highly aggressive tumors, including breast cancer. The association between MMP-2 and 9 and cell migration potential and invasiveness has been demonstrated (36). Furthermore, MMPs are released in an acidic environment, which activates them. The recruitment of numerous pH regulators, including NHEs, to the invadopodia membrane to control the surrounding microenvironment’s acidity and locally alkalinize the cytosol in breast cancer cells (37,38), underscores the potential impact of understanding these processes on cancer treatment.

In the present study, we described the inhibition of the Na^+^/H^+^ exchanger as an important mechanism of antitumor action of three copper(II) complexes containing 1,10-phenanthroline: [CuCl2(phen)·4H2O] (**1**), neocuproine (2,9 dimethyl-1,10 phenanthroline): [CuCl2(neo)·4H2O] (**2**) and 3,4,7,8 tetramethyl-1,10 phenanthroline: [CuCl2(tmp)·4H2O] (**3**) as the primary ligand and another three copper(II) complexes with the L-Ala-Phe dipeptide as an auxiliary ligand: [Cu(L-Ala-Phe)(phen)·4H2O] (**4**), [Cu(L-Ala-Phe)(neo)·4H2O] (**5**) and [Cu(L-Ala-Phe)(tmp)·4H2O] (**6**), on luminal subtype breast cancer cells (MCF-7 cell line) in the monolayer and spheroid type form.

## 2. Results and Discussion

### Effect of copper(II) complexes on cell viability and cell proliferation

Cytotoxicity studies for the six copper(II) complexes were performed through the MTT assay on MCF-7 (breast adenocarcinoma) and HaCaT (human keratinocyte) cells. To assess their antitumor effectiveness, we compared them with those of the reference metallodrug cisplatin. Figure 1, showed that **1** and **4** are the least effective in reducing cell viability, while the couples with neo and tmp showed 50% inhibition at 2.5 μM.

**Figure 1.**
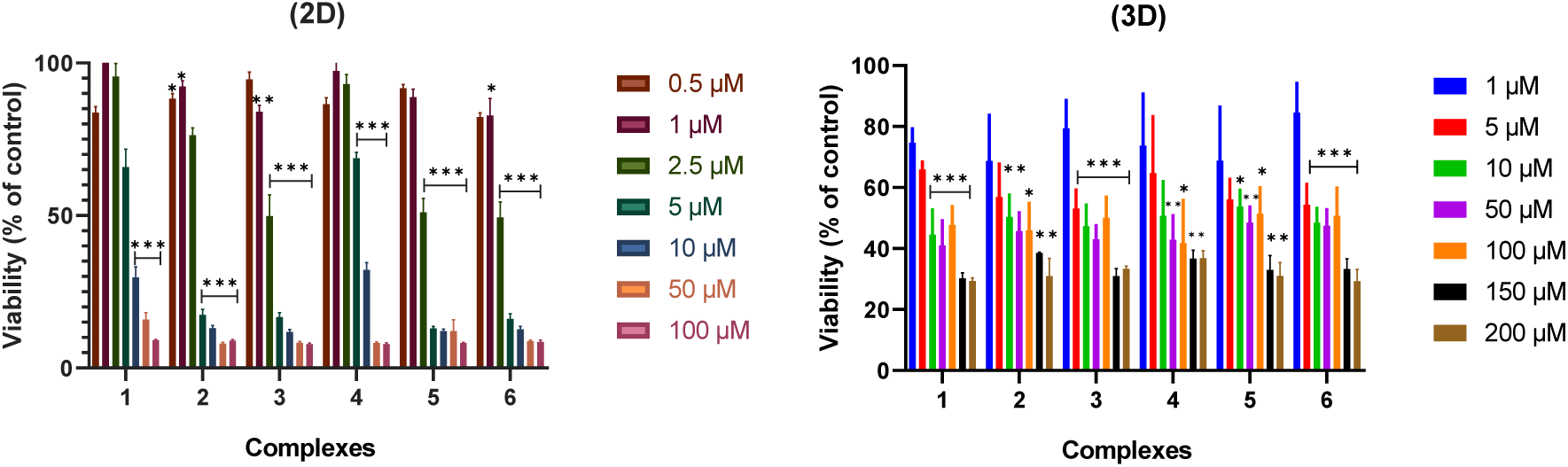
Cytotoxicity assay in 2D and 3D MCF-7 cell culture models with different concentrations of the complexes for 24 h. The results are expressed as the percentage of the basal level and represent the mean ± the standard error of the mean (SEM) (n=6). Asterisks represent a statistically significant difference in comparison with the basal level *(p<0.05) ** (p<0.01) ***(p<0.0001).

Table 1 shows that **3** exhibits a high cytotoxic activity after 24 h on MCF-7, displaying the lowest half maximal inhibitory concentration (IC_50_ = 1.8 μM), followed by **5** and **6** with IC_50_ = 2.2 μM . **1**, **2**, and **4** resulted in less activity, showing higher IC_50_ values. Nevertheless, all compounds are more active than cisplatin (IC_50_ = 19.6 μM in MCF-7) (39). The effect of the complexation was also established since the free ligand and the metal salt presented much higher IC_50_ values than all compounds tested. To understand the specific potential of the copper compounds and address their selectivity for cancer cells, we investigated their effect on the cell viability of human skin keratinocytes (HacaT cells). We compared their effects by calculating the selectivity index (SI). According to our findings, only the complexes with tmp (**3** and **6**) presented higher selectivity on MCF-7 cells than cisPt, with SI values of 1.3 and 2.2, respectively. In contrast, cisPt presents an SI value of 1.2 (the ratio of IC_50_HaCaT/IC_50_ MCF-7 calculates the SI value).

**Table 1.**
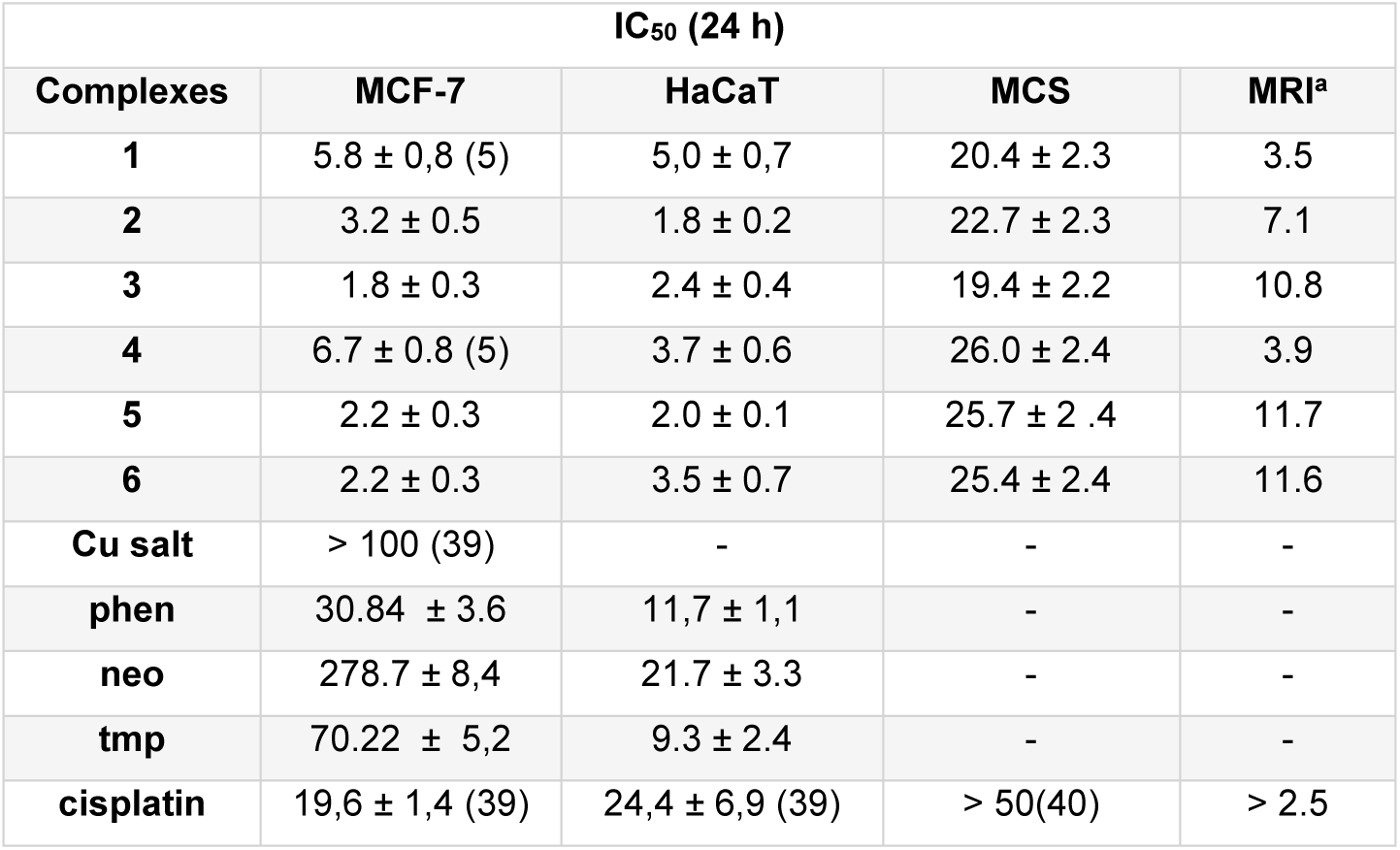
IC50 values of MCF-7 and HaCaT cells in the monolayer model and of MCF-7 cell in the multicellular spheroid (MCS) model for the six copper(II) complexes, a copper salt, phenanthroline, neocuproine, tetrametilphenantroline and cisplatin as positive control. ^a^ Multicellular Resistance Index (MRI) is the relationship between the IC50 obtained in spheroids divided by the IC50 obtained in the monolayer model.

On the other hand, the clonogenic assay was performed to evaluate the effect of the complexes on the cellular reproductive potential. Figure 2 shows a reduction of cell proliferation, which is in agreement with the cell viability assay on MCF-7 cells. In addition, all compounds affected the colony formation in a dose-dependent manner from 1.5 to 10 μM (all results under 50% survival). A total inhibition was observed at 3 μM for **6**, and at 5 μM for **3** and **5**. Again, **3** and **6** demonstrated a more significant inhibition effect than the rest.

**Figure 2.**
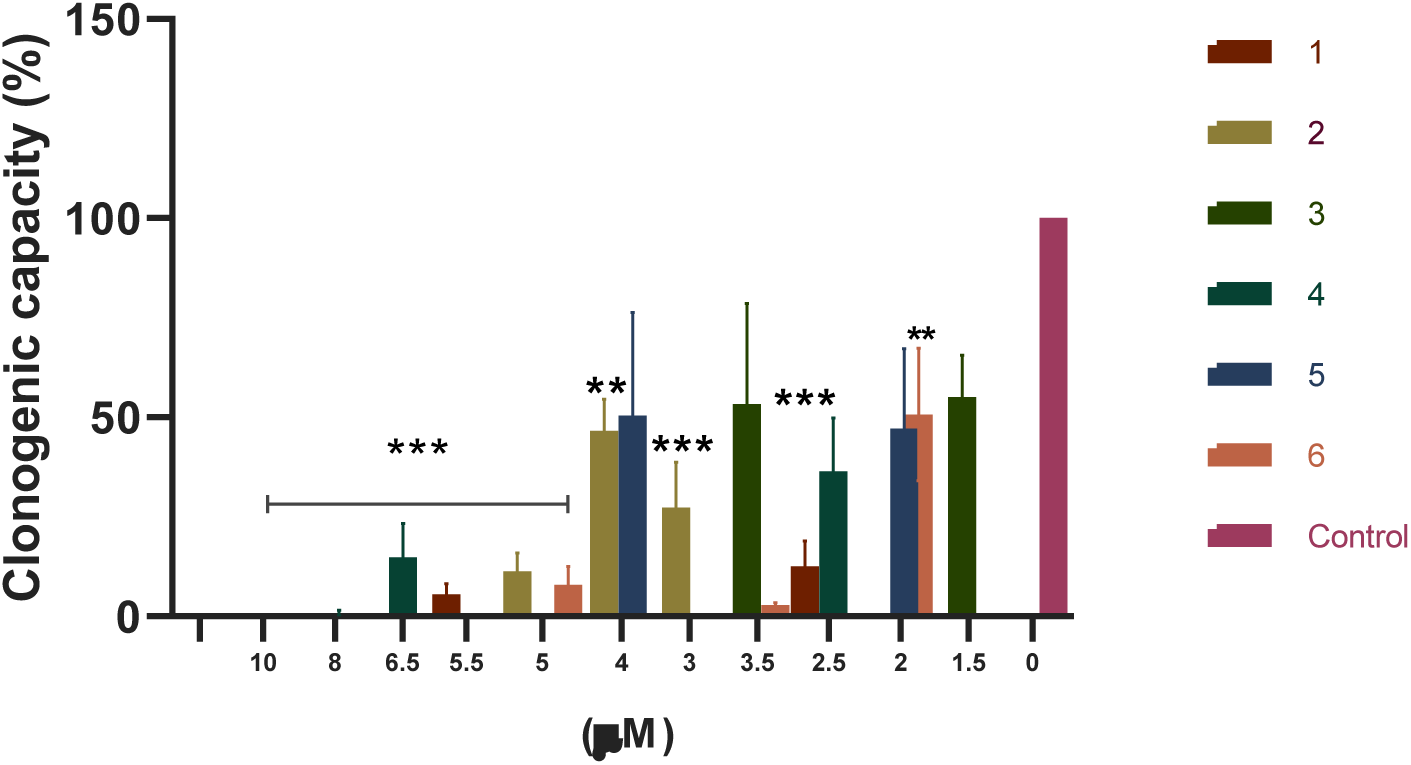
Clonogenic capacity in breast cancer cells (MCF-7) after treatment with different concentrations of the complexes at 37°C for 24h. The results are expressed as the surviving fraction as the percentage of the basal level and represent the mean ± SEM (n=4). Asterisks represent a statistically significant difference in comparison with the basal level **(p<0.05) ***(p<0.0001).

Different phenanthroline-based Cu(II) complexes were studied in various tumor cell lines, and similar results were obtained. It has been demonstrated that inserting a phenanthroline instead of a bipyridine-based ligand increased the antitumor effect in a series of Cu(II) proline-based complexes in A549 cells. Compounds with a substitution pattern in the fifth and sixth position of the phenanthroline backbone proved to be the most promising in terms of cytotoxicity (41) Heteroleptic copper(II) complexes having different phenanthroline derivatives showed antitumor action against breast cancer cells (MCF-7, MDA-MB-231, CAL-51) with IC_50_ values in the submicromolar range and outperformed clinically approved carboplatin. These complexes induced apoptotic cell death by inhibiting the proteasome (42).

Seng et al. showed selectivity towards cancerous nasopharyngeal HK1 cells rather than healthy NP69 ones when treated with different methylated glycine derivatives, with IC_50_ of 2.2 µM and Selectivity Index SI > 11.4 (43). Moreover, some of us synthesized ternary copper-dipeptide-neocuproine complexes and demonstrated that the introduction of neo as a ligand augmented the cytotoxic activity in different breast cancer cell lines after 48 h treatment (14).

### Effect of copper(II) complexes on oxidative stress

Our research on oxidative stress induction, a cell death mechanism that copper compounds could trigger, has been conducted with meticulous attention to detail. To evaluate the increase in ROS level, we assessed the impact of these complexes on the oxidation of DHR-123, a mitochondria-associated probe that selectively reacts with hydrogen peroxide, and DHE, which measures superoxide anion. This rigorous approach ensures the validity of our results and provides a solid foundation for further research in the mechanism of action.

Only 1 μM of **3** and **6** increased 320% and 137% the oxidation of DHE, respectively, and 1 μM of **6** significantly induced the oxidation of DHR-123 to 35% (Fig. 3). However, the impairment of the redox balance caused by **3** and **6** did not directly affect the cellular death process. Exogenous antioxidant scavenger NAC (N-Acetyl-L-Cysteine) or a mixture of vitamins C and E could not recover cell viability (not shown), indicating that the ROS production is unrelated to cell injury.

**Figure 3.**
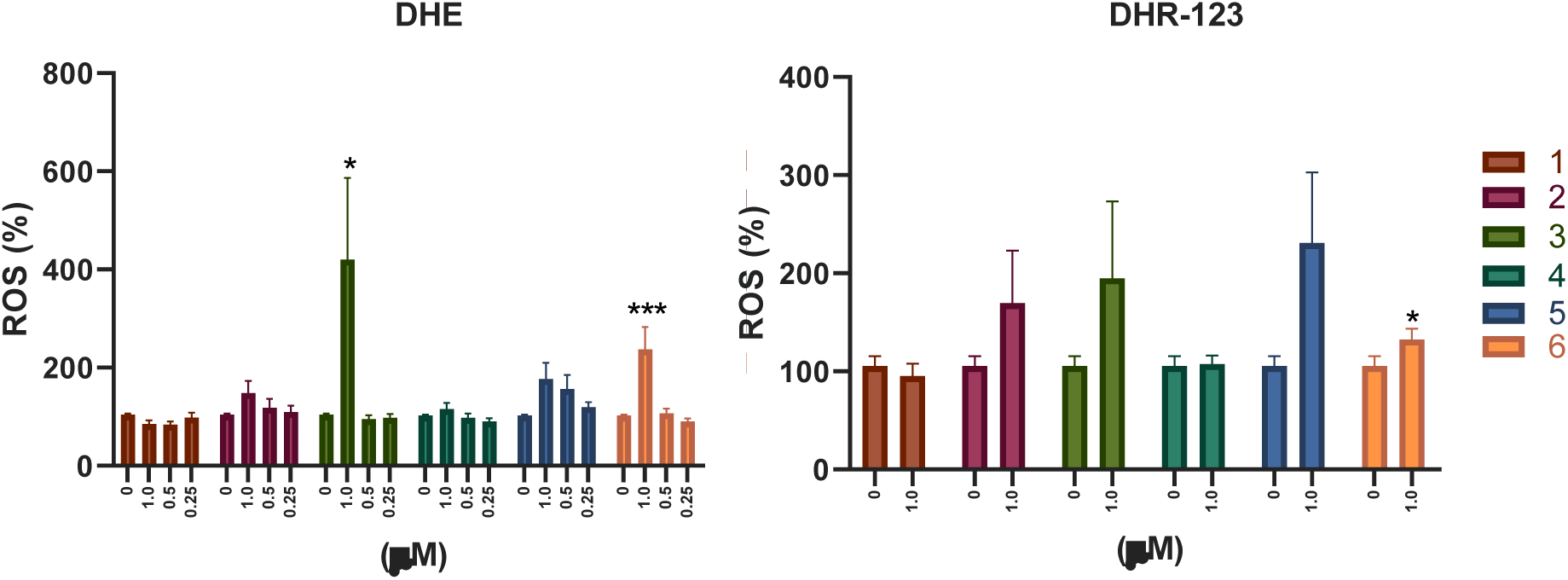
Induction of ROS by copper(II) complexes on MCF-7 cells by fluorometry (**a**) DHE and (**b**) DHR-123. The results are expressed as the mean ± SEM (n=3). * and *** represent a statistically significant difference in comparison with the basal level p < 0.05 and p < 0.001, respectively.

The copper(II)-phenanthroline complex with a glycine derivative, a subject of extensive research both in *vitro* and in *vivo* models, has shown significant potential in the fight against cancer. In agreement with our results, its anticancer activity is exerted through ROS accumulation, which causes mitochondrial dysfunction and DNA damage (44–46). Moreover, the copper(II) complex with phenanthroline and 5-hydroxytryptophan, shows no cytotoxic activity in normal phenotype MRC-5 cells but anticancer activity in A-549 cells in the micromolar range, under a mechanism based on cellular ROS production, GSH depletion and alteration of mitochondrial potential (47).

### Effect of copper(II) complexes on apoptosis induction

To study if the copper(II) complexes induce their cytotoxic effects through apoptotic processes, we measured phosphatidylserine’s externalization, a key event in the early stages of apoptosis, and cell integrity loss in the cell membrane using annexin V-FITC and propidium iodide. Two concentrations were employed to find apoptosis-related cell death, their IC_50_, and half the IC_50_. Only **3** induced late and early apoptosis, **4** and **6** induced early apoptosis in their IC_50_, and even more, **6** induced it at half the IC_50_ (Fig. 4; see results of **1** and **4** on (5), see the dot plot diagram of the flow cytometric analysis in Supplementary Material).

**Figure 4.**
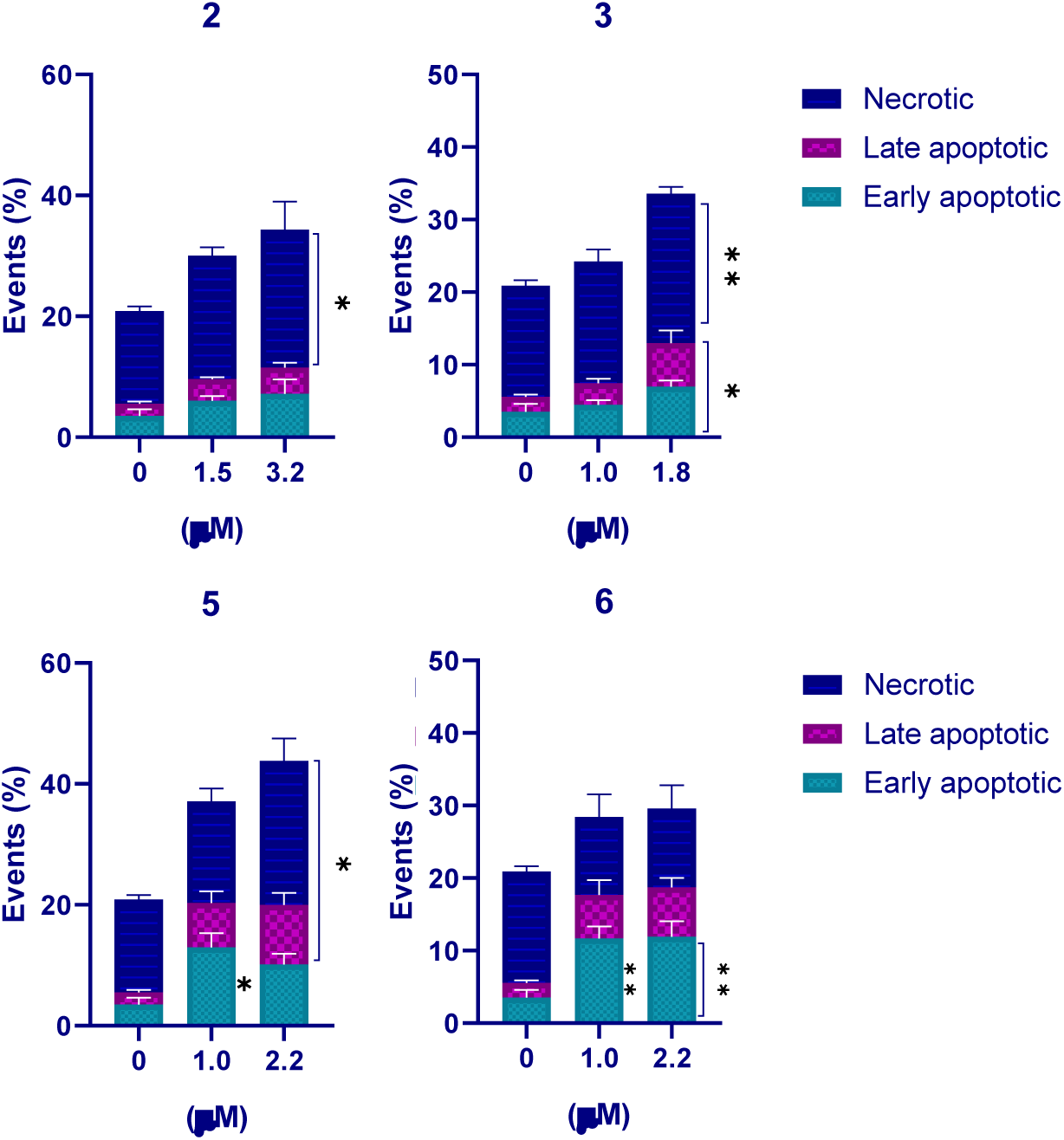
Determination of apoptosis by staining with Annexin V-FITC and Propidium iodide by flow cytometry in MCF-7 cells treated with the complexes for 24 h. The results are expressed as the average with the standard deviation of the error (SEM) (n=3). Asterisks indicate significant differences with the control * (p<0.05), ** (p<0.01), and *** (p<0.001). See the Supplementary Material for the Dot Plots).

Our results agree with the potential of two dinuclear copper(II) complexes with phenanthroline, which were cytotoxic in the micromolar concentration range towards cancerous A-549 and MCF-7 cancer cells and, to a lower extent, towards healthy HaCaT cells. Cell death resulted from an induction of the apoptotic pathway and increased G2/M cell cycle arrest. Interestingly, there is evidence of increased ROS production in A-549 cells. However, the copper complexes act as scavengers or inhibitors of ROS in the MCF-7 cell line as analyzed by DCFDA staining using flow cytometry (48).

### Effect of copper(II) complexes on genotoxicity

Genotoxicity induced by the copper complexes was studied by the alkaline comet assay, in which damaged DNA migrates out of the nucleus forming a tail, which can be quantified using image analysis. DNA damage was quantified as the comet tail moment, defined as the product of the tail length and the fraction of total DNA in the tail (Tail moment=tail length x % of DNA in the tail). MCF-7 cells were exposed to concentrations lower than the IC_50_ of all complexes (0.5 and 1 μM). Our results indicate that all compounds except **1** induced a genotoxic effect in MCF-7 cells at 1 μM (see results of **1** and **4** on (5), see representative pictures of comet induction in Supplementary Material). Moreover, **2** and **5** increased significant damage to DNA from 0.5 μM (see Fig. 5)(5).

**Figure 5.**
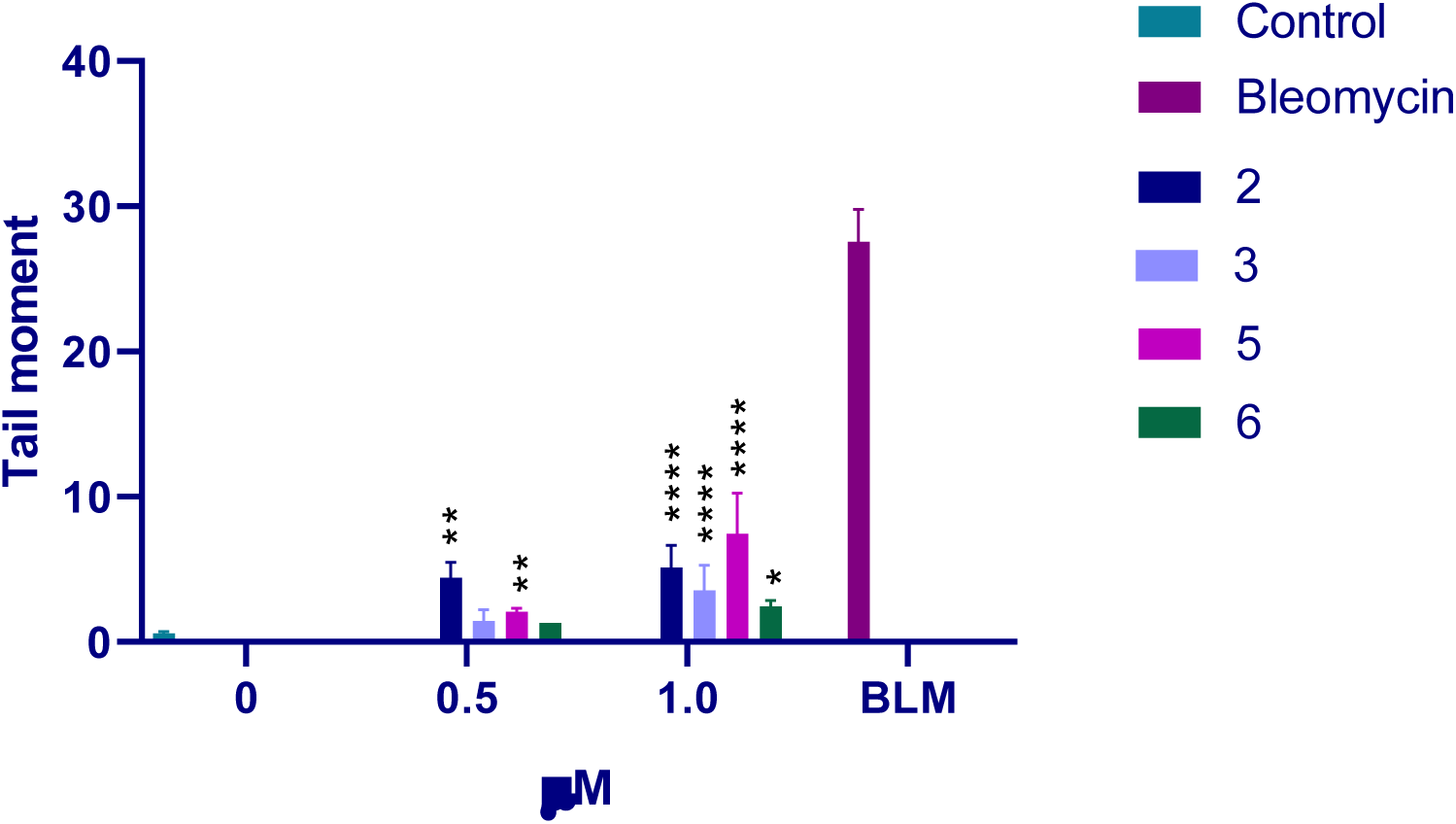
DNA damage (Tail moment) in MCF7 cells. The results are expressed as the average with SD (n=100). * (p<0.05), ** (p<0.01) and **** (p<0.0001)

Silva and coworkers (49) studied the interaction of two copper(II) compounds coordinated to two bidentate ligands, 1,10-phenanthroline and tetracycline or doxycyline. They have evaluated the binding of the complexes to DNA and their capacity to cleave it. The complexes bind to DNA preferentially by the major groove and then cleave its strands by an oxidative mechanism involving the generation of ROS (since radical inhibitors inhibit cleavage of DNA).

Some of us demonstrated earlier that copper(II) complexes with dipeptides as auxiliary ligands produce a hypochromic effect in the UV region observed upon the addition of CT-DNA. This binding event can be attributed to groove binding and or intercalation of the complex to the DNA (14). Other studies with ternary complexes with dipeptides showed anticancer potency in different cancer cell lines by cleaving at selective sites of G-quadruplex DNA; this meaningful interaction inhibits the telomerase, accumulating shorter telomeres and subsequent induction of apoptosis (50).

### Effect of copper(II) complexes on Na+/H+ exchanger-1 (NHE1)

Our study focuses on the effect of copper(II) complexes on the NHE1 activity, a pivotal player in tumor growth. NHE1 plays a central role in disrupting intracellular and extracellular pH homeostasis, promoting tumor cell migration and invasion, and, in some cases, conferring resistance to chemotherapeutic cell death (51). Despite its importance, research on the modulation of NHE1 by metal-based complexes remains limited. As expected, in non-treated cells, the NHE1 was hyperactivated whereas all the tested complexes were able to prevent its hyperactivity (Fig. 6) evidenced by the rate of intracellular pHi recovery from acidosis induced by the NH4^+^ prepulse technique. Interestingly, the effect was more pronounced in complexes **1**, **2**, **4**, and **5**, leading to intracellular acidification.

**Figure 6.**
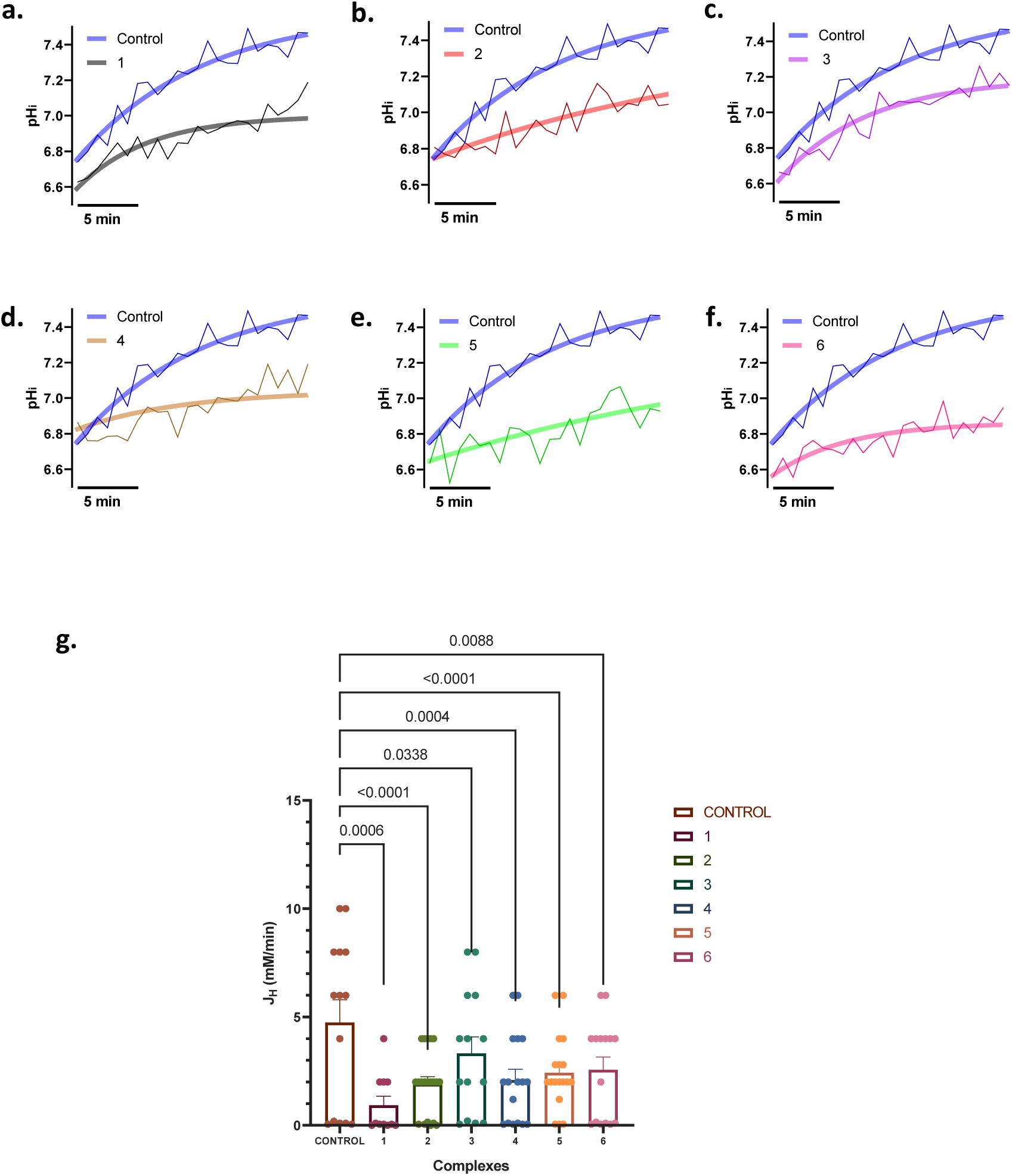
**a-f**. Representative traces of pHi recovery after the ammonium pulse (20mM NH4Cl) in MCF7 treated with the complexes and perfused with HK buffer. **g**. Average proton efflux (JH) carried by NHE, was calculated at different pHi values during the recovery from acidosis. The results are expressed as the percentage of the basal level and represent the mean ± the standard error of the mean (SEM) (n=3).

Cancer cells acidify the extracellular matrix with their high metabolic activity. Acid-extruding transports help further reduce the pH by adding protons taken out of the intracellular contents and left in the extracellular matrix. Studies have shown that knockdown or genetic deletion of such acid-extruding transports reduces tumor growth in several cancer models (52). The expression and activity of NHE are significantly increased in breast microinvasive foci and adjacent ductal cells in multicellular spheroids that resemble a ductal carcinoma *in situ*. This upregulation is closely associated with the local infiltration and distant metastasis of cancer cells, underscoring the gravity of the situation and the urgent need for further research in this area (53).

### Effect of copper(II) complexes on cell migration

Tumor cell migration is associated with liver, brain, and lung tumor metastasis. NHE1 could interfere with this process, as was observed in the study by Xiuju et al., where they removed the acid-extruding protein, and its absence reduced the rate of cell migration in MDA-MB-468 (54). In our study, the inhibition in cell migration was analyzed by the wound healing assay after a 24 h treatment. Our results (Fig. 7) showed that complexes **5** and **6** significantly inhibited cell migration in the studied tumor cell line as demonstrated statistically (See Supplementary Materials for results of the other complexes). Nevertheless, only **5** inhibited cell migration at the lowest concentration (0.5μM).

**Figure 7.**
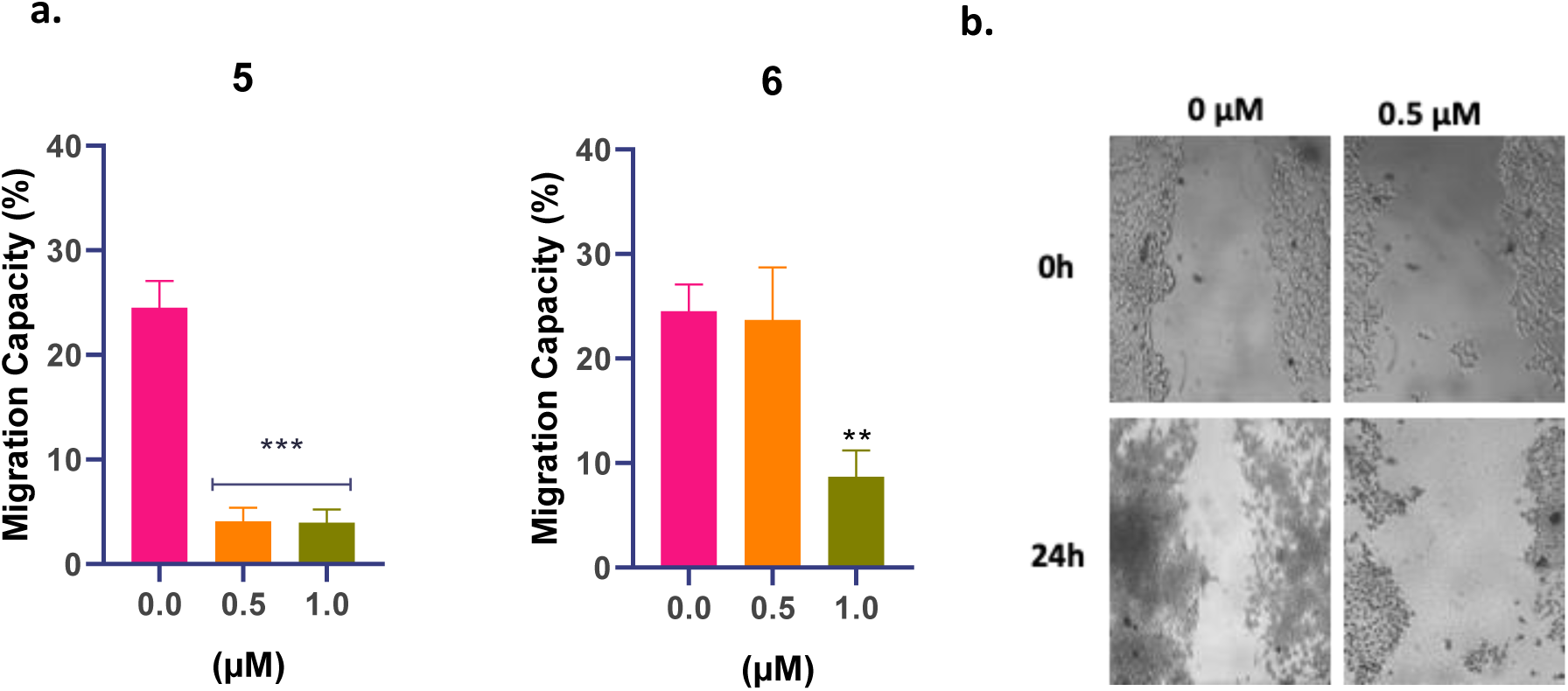
**a**. Cell migration assay in MCF-7 cells after 24 h treatment. The results are expressed as the average value with the standard deviation of the error (SEM) (n=3). Asterisks indicate significant differences with the control ** (p<0.01) and *** (p<0.001). **b.** Representative fields from one migration experiment at concentration 0.5 μM of **5**.

### Effect of copper(II) complexes on MMP-9 activity and expression

Matrix metalloproteinase-9 (MM-9) has an important role in processes such as inflammatory response, angiogenesis, wound healing, and the differentiation of human embryonic stem cells (55,56). However, high expression of MMP-9 increases tumorigenesis and metastasis(57,58). In particular, an elevated level of MMP-9 induces the metastasis and invasion of breast cancer, leading to death (59,60). Understanding the intricate role of MMP-9 in these processes is of huge importance, as it could improve novel therapeutic strategies in cancer treatment. In our study, complexes **4**, **5,** and **6** inhibited the activity MMP-9 (Fig. 8a) following the inhibition of Na^+^/H^+^ exchanger-1 (NHE1). Moreover, expression of MMP-2 and 9 measured by Western Blot, was also significantly affected by **1** and **4** (Fig. 8b). Previously, Yanxia Hu and coworkers observed that the activation of NHE1 can increase the activity of MMP-9 and MMP-2 and promotes invasive pseudopod maturation(61). In MDA-MB-231 breast cancer cells, NHE1 stimulated the expression of membrane type 1-MMP by activating the ERK1/2 and p38 MAPK signaling pathways by mediating the invasion and metastasis (62).

**Figure 8.**
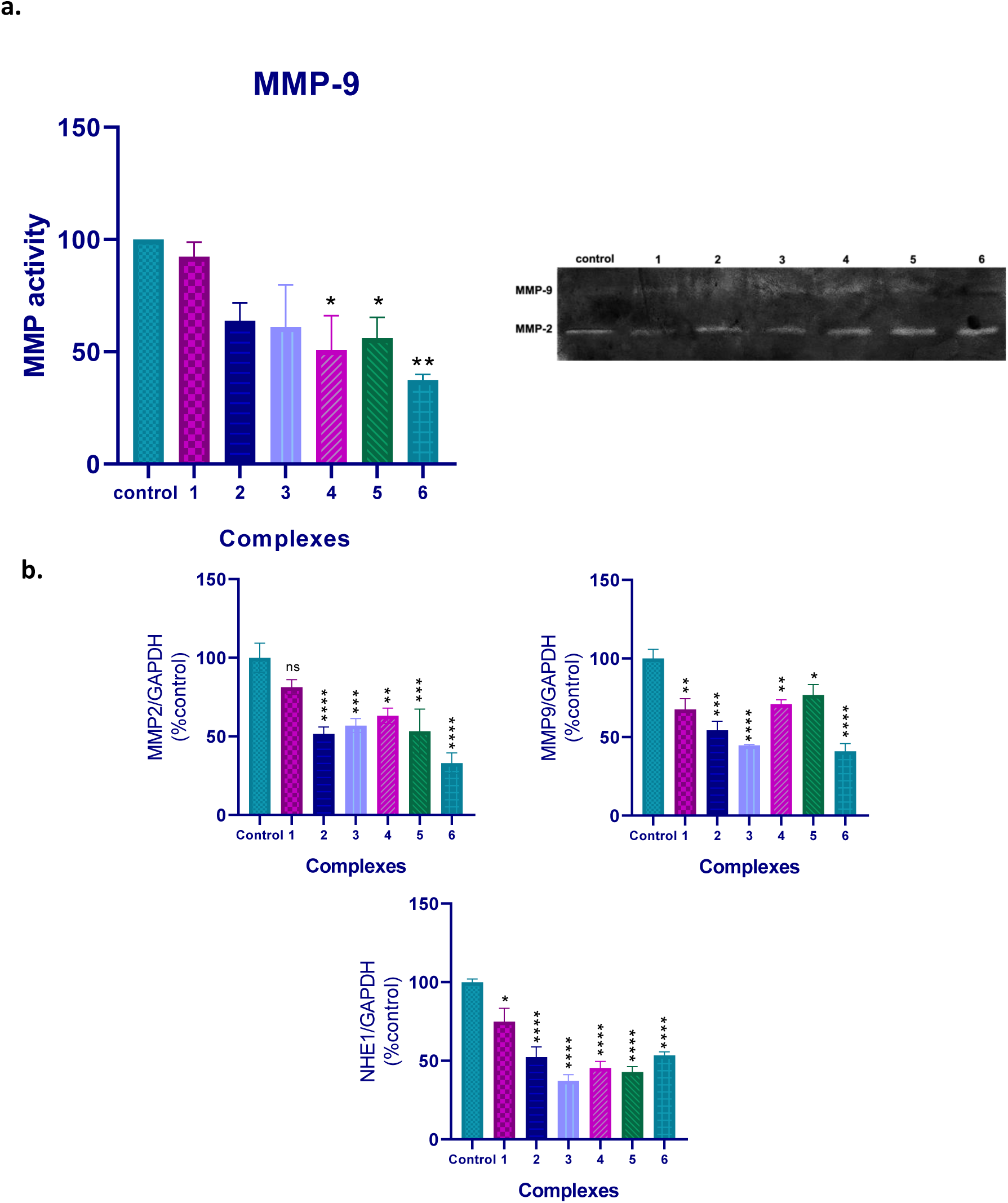
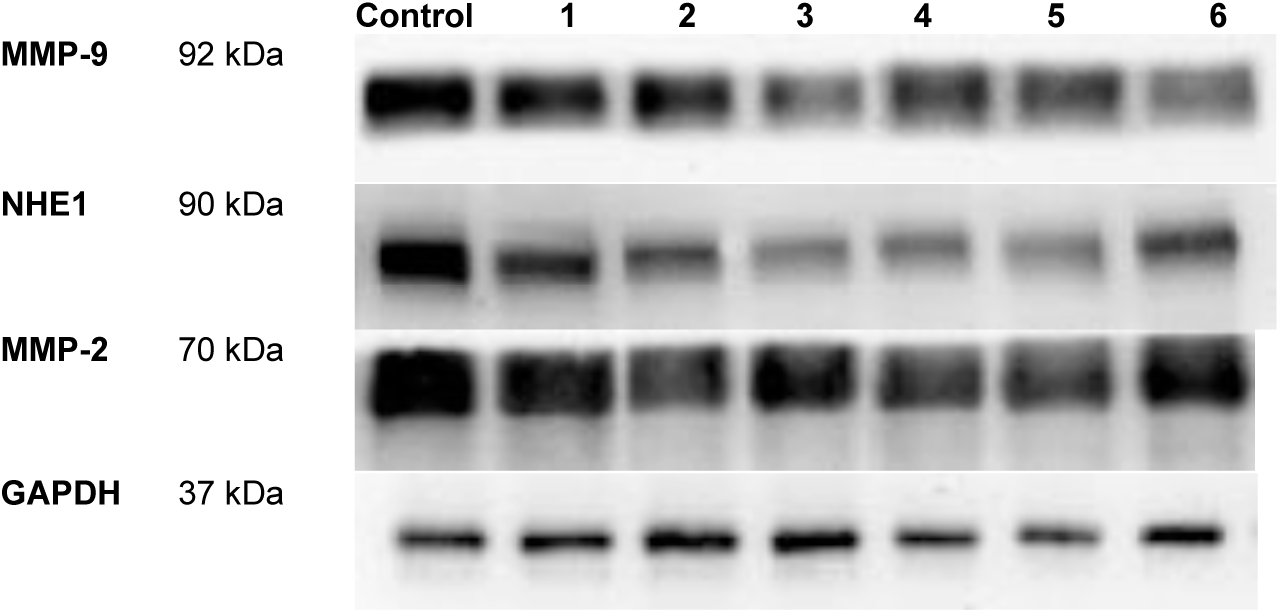
a. MMP-9 activity by zymography in MCF-7 extracts in 24 h. **b**. Western blot of lysates in the six Complexes for MMP-2, MMP-9 and NHE1. The results are expressed as the average with the standard deviation of the error (SEM) (n=3). Asterisks indicate significant difference with the control * (p<0.05), ** (p<0.01), *** (p<0.001)**** (p<0.0001).

Previous studies have demonstrated that paclitaxel-induced cell apoptosis is mediated by inhibiting NHE1 expression and activity. Furthermore, it has been confirmed that PKA and p38 are upstream regulatory points of NHE1. However, studies have also found that pyrazinoylguanidine-type NHE1 inhibitors potently inhibit the growth and survival of cancer cell spheroids, and this effect is unrelated to NHE1 inhibition. These findings could have significant implications for the development of novel cancer therapies (52).

### Effect of copper(II) complexes on multicellular spheroid (MCS) cell viability

Cell viability was evaluated on multicellular spheroids for the six complexes for 24 h. Our findings showed that copper complexes induced a concentration-dependent reduction in MCS viability from 1 μM, with a reduction of *ca.* 70% at 150 μM. Moreover, to cause the loss of viability of 50%, the concentration had to increase -compared to the one used in the monolayer model-several times (see IC_50_ from Table 1). Cisplatin, however, shows an IC_50_ value > 50 μM (40), at least twice higher than our complexes. Moreover, all complexes affected the shape and volume, and the edges became irregular on spheroids after 24 h. Our results are in agreement with copper(II) complexes tested in colorectal carcinoma cell spheroids, showing IC_50_ values from 15 to 30 μM after 48 h (63).

Multicellular spheroids are often employed as models to study tumor behavior because they better simulate the structure, microenvironment, and heterogeneity of actual tumors, which can affect drug resistance characteristics of solid tumors compared to traditional two-dimensional cell cultures. The Multicellular Resistance Index (MRI) measures the increased resistance of multicellular structures to treatments compared to single cells or 2D cell cultures, a fact observed for our compounds. The drug resistance mechanism can result due to cell quiescence, hypoxia-induced resistance, and drug penetration, since those therapeutic agents have difficulty penetrating the dense structure of the spheroid or because of higher expression of multidrug resistance transporters (e.g., ABC transporters), which pump drugs out of the cells, reducing their effectiveness (64,65).

## 3. Conclusions

Our study shows that copper(II)-phenanthroline-derivatives complexes exert a better antitumor effect than the reference drug cisplatin, currently used in the clinic for different solid tumors, in both models of breast cancer cells.

However, we found discrepancies between them: the pair containing the tmp as the primary ligand improves the antitumor activity demonstrated by lower IC_50_ values, higher selectivity indexes, apoptosis induction, and better inhibition of cell proliferation. Furthermore, the primary ligand is important in generating reactive oxygen species in this cell line. However, this imbalance in the redox status is unrelated to cytotoxicity.

Along with the cytotoxicity, all copper complexes promoted DNA damage, as demonstrated by the comet assay. Moreover, copper(II) complexes inhibited the expression and activity of NHE1 exchanger in MCF-7 cells, and the expression of MMP2, MMP9. However, only dipeptide-containing complexes showed a decrease in MMP-9 activity and an inhibition of cell migration.

## 4. Acknowledgements

This work was supported by UNLP (2020/2023 X899), CONICET (PIP 0235) and ANPCyT (PICT 2021-00090, PICT 2021-00338) from Argentina.

## 6. Experimental section

### Materials

Tissue culture materials were purchased from Corning (Princeton, NJ, EUA) and APBiotech (Buenos Aires, Argentina). Dulbecco’s Modified Eagle Medium (DMEM) and TrypLE™ from Gibco (Gaithersburg, MD, EUA) and fetal bovine serum (FBS) from Internegocios S.A. (Buenos Aires, Argentina); Rhodamine 123, dihydroethidium (DHE), annexin V-FITC, propidium iodide (PI), SYBR Green, low melting point agarose from Invitrogen Corporation (Buenos Aires, Argentina). The vitamin E was purchased from Sigma Aldrich (St. Louis, MO, EE. UU.) and vitamin C from Merck (Buenos Aires, Argentina). Bleomycin (BLM) (Blocamycin) was kindly provided by Gador S.A. (Buenos Aires, Argentina). MCF-7 (HTB-22) and HACAT cell lines were purchased from ATCC ®, PVDF membrane (BioRad, CA, USA), BCECF-AM (Thermofisfher, USA), APH (sigma).

### Synthesis of copper complexes and aqueous stability

Six copper (II) complexes [CuCl_2_(phen)] (**1**), [CuCl_2_(neo)] (**2**), [CuCl_2_(tmp)] (**3**), [Cu(L-Ala-Phe)(phen)·4H_2_O] (**4**), [Cu(L-Ala-Phe)(neo)·4H_2_O] (**5**) and [Cu(L-Ala-Phe)(tmp)·4H_2_O] (**6**) where phen stands for 1,10-phenanthroline, neo for neocuproine (2,9-dimethyl-1,10-phenanthroline) and tmp for 3,4,7,8-tetramethyl-1,10-phenanthroline, were synthesized and physicochemical characterized at the Faculty of Chemistry, University of the Republic, Montevideo, Uruguay. The purity of the samples used in this study was confirmed by elemental analysis by comparison with recent reports for both compounds (66) Experimental elemental analysis for (1): Calc. for CuCl_2_C_12_H_9_N_2_O 0,5 PM 323,66: found %C 44.53/44.11, %N 8.66/8.36, %H_2_.80/2.92 y %S 0.00/0.00. (2): Calc. for CuCl_2_C_14_H_13_,5N_2_O0.75 /Found %C: 47.20/47.16 %N: 7.86/7.62 %H: 3.82/3.86. (3): Calc. for C_16_H_18_CuN_2_OCl_2_/Found: %C: 7.21/7.08, %N: 49.43/49.09, %H: 4.67/4.61. (4): Calc. for C_24_H_30_CuN_4_O7: C, 52.40, N, 10.19, H, 5.50 found: C, 51.85, N, 10.17, H, 5.12. (5): Calc. for C_24_H_30_CuN_4_O_7_: C, 52.40, N, 10.19, H, 5.50 Found: C, 52.05, N, 10.37, H, 5.15. (6): Calc./Found (C_28_H_36_CuN_4_O_6_) %C: 57.18/57.52, %N: 9.52/9.40, %H: 6.17/6.07.

Fresh stock solutions of **1**, **2**, **4**, and **6** (10 mM) were prepared in PBS and diluted with culture medium according to the concentrations indicated in the results section. Fresh stock solutions of **3** and **5** (10 mM) were prepared in DMSO and diluted with culture medium according to the concentrations indicated in the figures. We used 0.5% as the maximum DMSO concentration to avoid the toxic effects of this solvent on the cells (5,14,15).

### Methods Cell culture

Human breast cancer line (MCF-7) and Human keratinocyte cell line (HaCaT) were cultured in DMEM medium supplemented with 10% fetal bovine serum (FBS), 100U/ml penicillin, and 100 μg/ml streptomycin at 37°C in a humidified atmosphere with 5% CO_2_. These cell lines were seeded in a T75 flask and when they reached 80-90% of confluence, cells were subculture using TrypLETM. Experiments were carried out in multiwell culture plates where cells were allowed to attach and treated for each complex.

### Cytotoxicity assay

Cell viability was determined by the 3-(4,5-dimethylthiazol-2-yl)-2,5-diphenyltetrazolium bromide (MTT) method, which is reduced by mitochondrial dehydrogenases in viable cells to a purple formazan(67). Briefly, 1.5x10^4^ cells were seeded on 96 well-cultured plates and incubated at 37 °C in a humidified atmosphere with 5% CO_2_. After 24 h, cells were exposed to different concentrations of each complex, metallic salt, and ligand (in the concentration range from 0.5-100 μM) for 24 h. Afterward, the monolayers were washed and incubated with 0.5 mg/mL MTT in DMEM for 3 h. Color development was measured spectrophotometrically with a microplate reader (multiplate reader Multiskan FC, Thermo Scientific) at 570 nm after dissolving the formazan precipitate in DMSO (100 µL per well). Cell viability was plotted as the percentage of the control value. Additionally, cells were incubated with 10 μM Cariporide (an NHE1 inhibitor) to verify that cell death occurs due to the inhibition of NHE1 exchanger.

A mixture of vitamins C and E (50 μM each) or NAC was added to the culture medium with the complexes to evaluate the role of oxidative stress in the cytotoxicity. In the case of NAC, the monolayer was pre-treated with 250 μM NAC for 2h, and the culture medium was replaced with concentrations of the complexes. In both cases, cytotoxicity was evaluated using the MTT method.

### Clonogenicity method

A clonogenic assay was conducted according to Puck and Marcos (68) with minor modifications to explore whether the compounds affect cell proliferation. Cells were seeded and treated with different concentrations of each complex for 24h. After this time, monolayers were harvested with TrypLE™ and manually counted after dying the living ones with trypan blue. The cells were diluted to 10^3^ cells per well on a 12-well plate. After incubation for 7 days, cells were washed with PBS and stained with 0.5% crystal violet for 30 min at room temperature. Each colony consists of at least 20 cells. Colonies were counted under an Olympus BX51 inverted microscope.

### Determination of reactive oxygen species (ROS) production

ROS induction in MCF-7 cells was measured as a mechanism of cell death. Briefly, cells were seeded in 24-well plates, allowed to attach for 24 h, and treated with different concentrations of complexes for 24 h at 37°C. Afterward, the monolayers were washed with Hank’s solution (NaCl 0.137 M, KCl 0.8 mM, MgSO_4_.7H_2_O 0.8 mM, CaCl.H_2_O 1 mM, KH_2_PO_4_ 0.4 mM, C_6_H_12_O_6_ 5 mM, NaHCO_3_ 4 mM, Na_2_HPO_4_ 0.3 mM), added 480 μl DHR-123 or DHE, incubated for 30 min at 37°C in the darkness. Cells were rewashed with Hank’s solution and incubated for 45 min at 37°C with lysis solution (Triton X-100 0.1%). ROS generation was determined by the oxidation of DHR-123 to rhodamine or by the interaction of DHE and superoxide ion (O2-^—^) (69). Fluorescence was registered using a fluorometer Shimadzu RF-6000 (λ excitation: 485 nm; λ emission: 530 nm, for DHR-123; and λ excitation: 500 nm; λ emission: 580 nm, for DHE). H_2_O_2_ 300 mM for 20 min was employed as a positive control. Results were corrected for protein content, measured with the bicinchoninic acid (BCA) method.

### Bicinchoninic acid assay

The bicinchoninic acid assay determined protein content according to Smith et al., (70) in which proteins reduce Cu^2+^ to Cu^+^ in an alkaline medium (Biuret reaction). Proteins react with BCA to produce a deep violet color measured at 570 nm. Briefly, 25 μl of the cell extract described above, was incubated with 200 μl of the BCA standard for 30 min at 37°C. The BCA-Copper complex reaction was analyzed by measuring absorbance on a Multiskan FC Thermo Fisher plate reader.

### Apoptosis

Cell death induction was evaluated with Annexin V-FITC and propidium iodide (PI) staining. Briefly, MCF-7 cells were seeded in 12-well plates and incubated overnight. Copper(II) complexes were added at the indicated concentrations for 24 h. After treatment, cells were washed with PBS, treated with trypsin, and centrifuged. The pellets were resuspended in 300 μl of binding buffer with 1 μl de annexin V-FITC and 1μl de PI. Cells were analyzed by flow cytometry with a Becton Dickinson FACScaliburTM flow cytometer (BD Biosciences, USA) and further analyses were performed using Flowing 2.5 software. 10^4^ events were acquired per sample.

### Single-cell gel electrophoresis assay

To detect DNA damage, the single-cell gel electrophoresis assay (Comet assay) was employed based on the method of Singh et al. (71) with minor modifications. Briefly, MCF-7 cells were treated with different concentrations of the complexes. After 24 h, cells were suspended in 0.5% low melting point agarose and immediately poured onto microscope slides precoated with 0.5% normal melting point agarose. Slides were immersed in ice-cold lysis solution (2.5 M NaCl, 100 mM Na_2_-EDTA, 10 mM Tris–HCl, 1% Triton X-100, 10% DMSO at 4 °C, pH 10) for 1 h. Then, the slides were placed on a horizontal gel electrophoresis tank, and the DNA was allowed to unwind for 20 min in freshly prepared alkaline electrophoresis buffer (300 mM NaOH and 1 mM Na_2_-EDTA, pH 12.7). Electrophoresis was carried out in the same buffer for 30 min at 25 V (≈0.8 V/ cm across the gels and ≈ 300 mA) in an ice bath condition. Afterward, the slides were neutralized and stained with SYBR Green. The analysis was performed in a Nikon eclipse e200 fluorescence microscope. One hundred randomly captured cells per experimental point were used to determine the tail moment using Comet Score version 1.5 software. Just before the cells were harvested, a pulse of 20 min of 10 μg/mL bleomycin just before the cells were harvested was employed as the positive control.

### pH measurement and ammonium pulse

The NHE1 activity was assessed by evaluating the pHi recovery from an ammonium pre-pulse-induced acute acid load. Intracellular pH (pHi) was measured by BCECF-AM (2′,7′-Bis-(2-Carboxyethyl)-5-(and-6)-Carboxyfluorescein, Acetoxymethyl Ester)-epifluorescence technique (71). Briefly, MCF-7 cells were seeded in 48-well plates and incubated overnight. Copper(II) complexes were added at the indicated concentrations for 24 h. After treatment, cells were incubated for 15 min at 37°C with 10 μM BCECF-AM dissolved in Krebs-Henseleit (KH) solution (in mM): 146.2 NaCl, 4.7 KCl, 1 CaCl2, 10 HEPES, 0.35 NaH2PO4, 1 MgSO4, and 10 glucose (pH adjusted to 7.4 with NaOH). Steady-state pH was measured for 10 min in KH, next transient exposure to 20 mM NH4Cl was made for 10 min and finally, the recovery was recorded for 15 min in KH. BCECF-AM fluorescence was measured through dual excitation (440 and 485 nm) and emission at 520 nm in a spectrofluorometer plate reader Varioskan LUX (Thermofisher). The 495-to-440 nm fluorescence ratio was calculated, and, at the end of each experiment, the fluorescence ratio was converted to pH by calibrations using the high K^+^-nigericin method (72). Proton efflux (JH) was calculated as dpHi/dt x βi at a common pHi of 6.8. The dpHi/dt at each pHi was obtained from an exponential fit of the recovery phase. βi is the intracellular buffering capacity and was measured by exposing MCF-7 cells to varying concentrations of NH4Cl in 1Na^+^-free KH solution, next, βi= Δ[NH4^+^]i/ΔpHi and referred to the mid-point values of the measured changes in pHi. βi at different levels of pHi were estimated from the least squares regression lines βi vs. pHi plots(73).

### Wound healing assay

MCF-7 cells were grown in 12-well culture plates with DMEM medium supplemented with 10% (FBS) to achieve an 80-90% confluent monolayer. After this, a superficial simple scratch wound was created in the monolayer, washed with PBS, and cells were treated for 24 h with 1.0 and 0.5 μM copper(II) complexes. The monolayer was washed with PBS and stained with Giemsa; the wound was photographed using an Olympus BX51 inverted microscope with a digital camera. Cell migration was analyzed with ImageJ Software. The percentage of migration was calculated using the following formula: (final area (TF) – initial area (T0) / initial area (T0).

### Gelatin Zymography assay

The Gelatin Zymography assay, a practical method based on the work method of Rajeev B et al. (72), is a useful tool to analyze the activity of metalloproteinases (MPPs). Cells were seeded in 12-well plates containing DMEM medium supplemented with 10% of FBS, and treated with the complexes for 24h. Cells were incubated at 37°C with serum-free culture media overnight. The cell-conditioned media were then electrophoresed in a 10% acrylamide gel containing 0.1% gelatin. After electrophoresis, the gel was incubated at 37°C for 48h in a buffer (50 mM Tris, 200 mM NaCl, 5 mM CaCl_2_, triton, pH 7.4) to facilitate gelatin degradation. The gel was stained with 0.5% Coomassie Brilliant Blue and discolored in a destaining solution. Bands were quantified using Image J and relativized to total protein.

### Western blotting

To determine the expression of metalloproteinases (MMPs) and NHE1, cells were seeded in 6 wells plates overnight, and treated with the complexes for 24 h. Next, cells were lysed by adding cold RIPA buffer (KH_2_PO_4_ 30 mM, NaF 25 mM, EDTA 5 mM, sacarose 300 mM, Triton X-100 0.01 %, Igepal 1 %) with a protease inhibitor cocktail (Roche). 60 ug of protein (measured by the Bradford method (75)) were loaded with 8% SDS polyacrylamide separating gel and transferred to polyvinylidene difluoride membranes (PVDF) membranes. Each membrane was incubated with the specific primary antibodies: NHE-1, MMP2, MMP9, GAPDH, as internal control (1:1000; Santa Cruz Biotechnology); and secondary antibodies: anti-mouse (1:10000; Cell Signaling). Immunoreactivity was visualized by a peroxidase-based chemiluminescence detection kit (Immobilon Western Millipore) using a Chemidoc Imaging System. The signal intensity of the bands in the immunoblots was quantified by densitometry using Image J software (NIH, USA).

### Spheroids formation

Spheroids of MCF-7 cells were developed and cultivated in 96-well plates with 100 µL of agar 1% and DMEM with glutamine without FBS. After five days of culture, the morphology was observed in a Microscope Nikon Eclipse TE2000-S. The MCF-7 cells-derived spheroids were treated at the indicated concentrations of the complexes for 24 h. Spheroids were centrifuged twice and washed with PBS. Afterward, spheroids were incubated with acid phosphatase buffer (APH) at 37°C for 3 h and revealed with 10 μl NaOH 1N. The analysis was performed in a multi-mode microplate reader SLXFATS.

### Statistical analysis

Results are expressed as the mean of independent experiments and represented as mean ± standard error of the mean (SEM). The total number of repeats (n) is specified in the legends of the figures. The Tukey test (two-way ANOVA) was employed to compare means in all the experiments.

## Supplementary Material

**A.** Dot Plot diagram of the flow cytometric analysis of MCF-7 cells treated for 24 h with all complexes. The X axis (FL1) corresponds to labeling with propidium iodide and the Y axis (FL2) relates to the cell population labeled with Annexin V-FITC. **1** and **4** were described at Munoz (5).

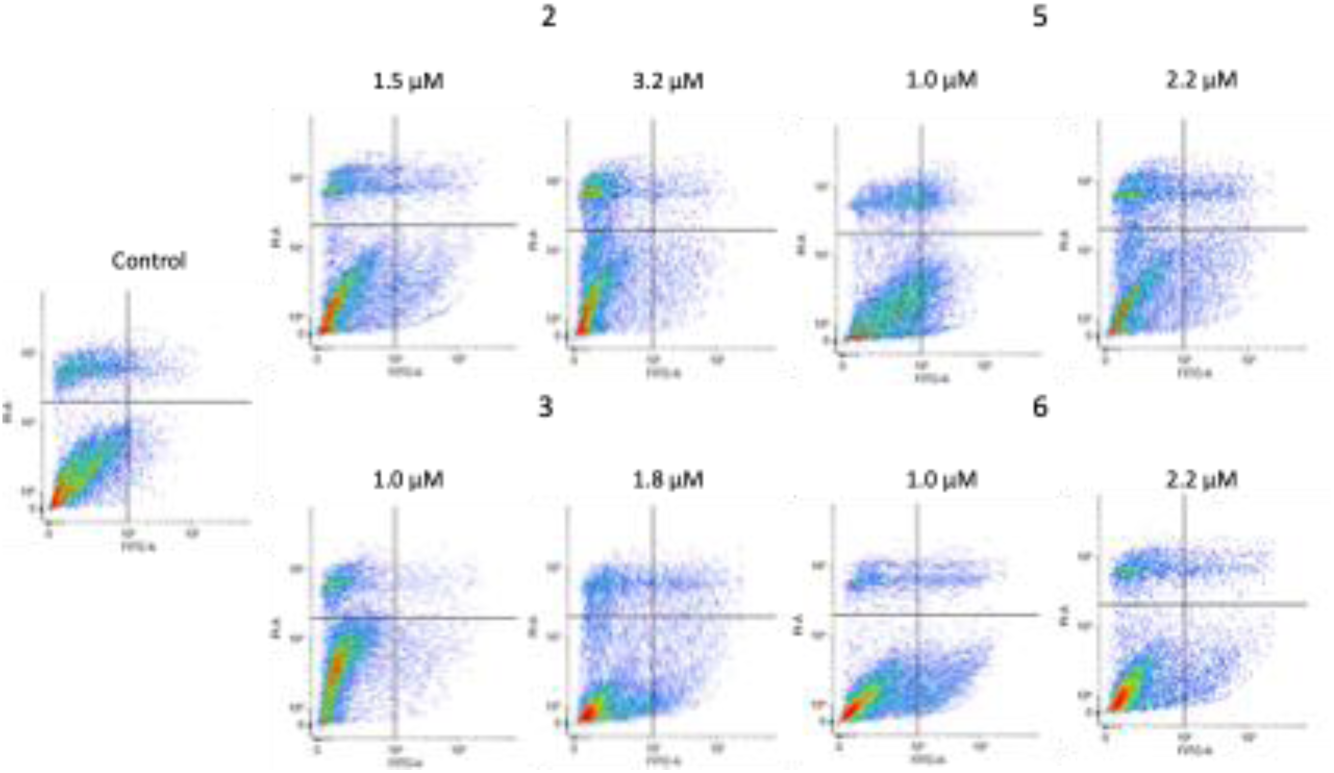

**B.** Induction of genotoxicity in complexes **2**, **3**, **5** and **6** in MCF-7 cells by comet assay. **1**

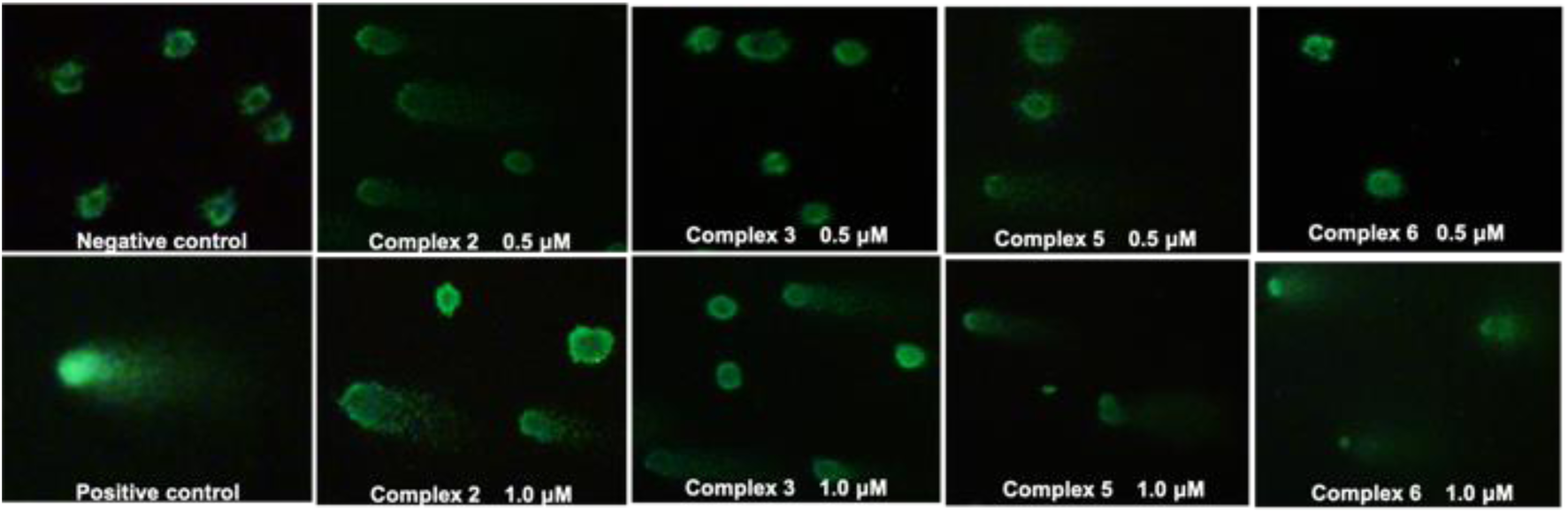

and **4** were described at Muñoz et al. (5).

**C.** Cell migration assay in MCF-7 for **1**,**2**,**3** and **4**. The decrease in migration capacity was not significantly different.

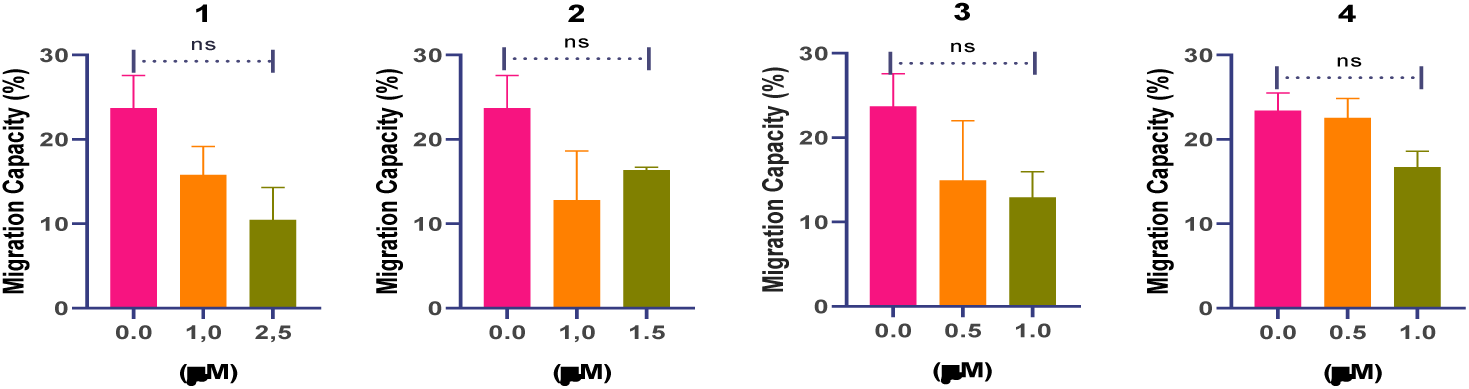

